# Nucleotide resolution genetic mapping in pigs by publicly accessible whole genome imputation

**DOI:** 10.1101/2022.05.18.492518

**Authors:** Rongrong Ding, Rodrigo Savegnago, Jinding Liu, Nanye Long, Cheng Tan, Gengyuan Cai, Zhanwei Zhuang, Jie Wu, Ming Yang, Yibin Qiu, Donglin Ruan, Jianping Quan, Enqin Zheng, Linjun Hong, Zicong Li, Suxu Tan, Mohammd Bedhane, Robert Schnabel, Juan Steibel, Cedric Gondro, Jie Yang, Wen Huang, Zhenfang Wu

## Abstract

Genetic mapping to identify genes and alleles associated with or causing economically important quantitative trait variation in livestock animals such as pigs is a major goal in the genetic improvement animals. Despite recent advances in high throughput genotyping technologies, resolution of genetic mapping in pigs remains poor due in part to the low density of genotyped variant sites. In this study, we overcame this limitation by developing a reference haplotype panel for pigs based on 2,259 whole genome sequenced animals representing 44 pig breeds. We optimized the imputation procedure to achieve an average concordance rate in excess of 97%, non-reference concordance rate 91%, and *r*^2^ 0.89. We demonstrated that genotype imputation using this resource can dramatically improve resolution of genetic mapping. Finally, we developed a public web server (swimgeno.org) to allow the pig genetics community to fully utilize this resource. We expect the resource and server to significantly facilitate genetic mapping and accelerate genetic improvement in pigs.

## Introduction

The domestic pig (*Sus scrofa*) is an important livestock species and a model organism for biomedical research (Lunney et al., 2021). Historically, domestication and intense artificial selection have created many pig breeds that are genetically and phenotypically distinct from each other and from their wild relatives (Bosse et al., 2014; Groenen et al., 2012; Li et al., 2013). More recently, high throughput DNA sequencing and genotyping technologies (Ramos et al., 2009) have facilitated the genetic improvement of pigs. For example, hundreds of genome-wide association and quantitative trait locus (QTL) mapping studies have identified numerous genomic regions associated with various production, physiological, and behavioral phenotypes (Hu et al., 2019). These studies are important for understanding the genetic and biological basis of economically and biomedically important traits such as growth (Onteru et al., 2013), fertility (Sell-Kubiak et al., 2015), and disease resistance (Boddicker et al., 2014).

The resolution of genetic mapping in pigs remains poor, due in part to the low density of single nucleotide polymorphism (SNP) genotyping arrays. One proven cost-effective approach to overcome the limitation in resolution is through genotype imputation, leveraging linkage disequilibrium to infer genotypes at unobserved polymorphic loci (Marchini and Howie, 2010). With large haplotype reference panels created by whole genome sequencing, imputation has the potential to provide sequence level genotypes (Das et al., 2016). In livestock animals, where QTL identification and genetic prediction are two major goals and linkage disequilibrium is extensive, sequence level genotype imputation has been successfully applied with a relatively small number of reference haplotypes but decent accuracy (Van Den Berg et al., 2019; Daetwyler et al., 2014). However, there is a lack of easily accessible public resources to deliver high accuracy imputed genotypes from diverse reference populations, as exemplified by the public Michigan Imputation Server for humans (Das et al., 2016).

In this study, we produced whole genome sequence data from 1,530 newly sequenced pigs and combined them with 729 additional animals from public databases to call variants and develop by far the largest and most diverse reference panel of haplotypes in pigs to date. This substantial increase in the number of available genomes allowed us to impute SNP array genotypes to whole genome sequences rapidly and accurately. We evaluated accuracy of imputation and demonstrated the utility of this haplotype reference panel in genome-wide association mapping. We introduce a new public web server (swimgeno.org) where users may submit array genotypes and retrieve imputed whole genome sequence level genotypes. This resource will greatly improve accessibility to high accuracy genotype imputation, facilitating potentially nucleotide resolution genetic mapping in pigs.

## Results

### Patterns of genetic variation in >2,000 pig genomes

We consolidated whole-genome sequence data from newly sequenced animals (*n* = 1,530) and publicly available data (Table S1-2) for a total of 2,259 pigs, representing 44 different breeds (Table S1). The majority of animals were Landrace (*n* = 651), Yorkshire (*n* = 543), and Duroc (*n* = 485), three major commercial breeds. The uniquely aligned sequence depth was approximately 12.86 X averaged across all animals (Table S1). We called variants using the GATK pipeline and calibrated variant quality scores with known variant sets compiled from commercial SNP arrays. After filtering out variants of low quality and excessive heterozygosity and missingness, 47.86 M autosomal variants remained. Sub-sampling of animals indicated that the increase in the number of discovered variants quickly diminished (Figure 1a). More than 95% of all variants could be recovered using only 1,000 randomly selected animals, suggesting that this dataset likely captured the vast majority of segregating variants in the global pig population.

**Figure 1.**
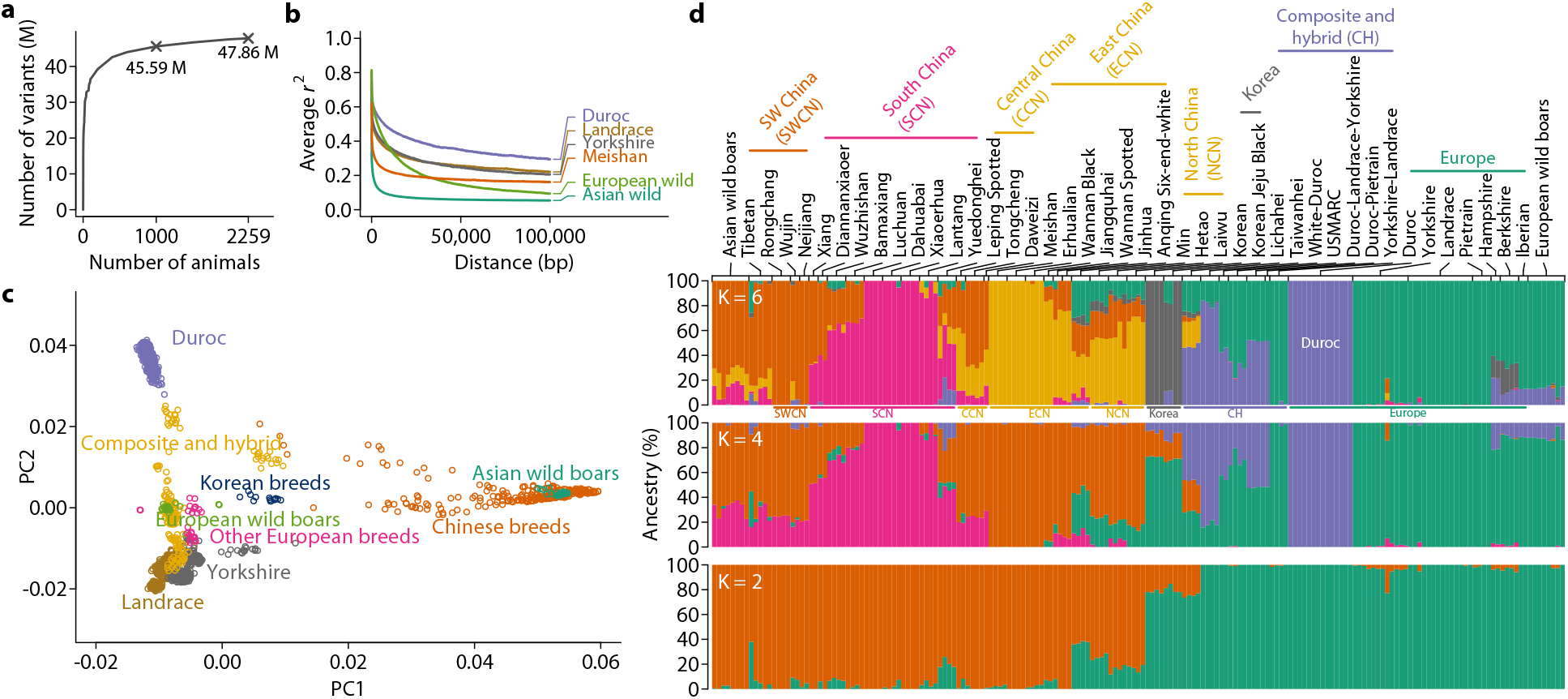
Genetic structure of the global pig population. (**a**) Number of variants discovered as a function of number of animals in the discovery cohort. The curve is generated by randomly subsetting the population and counting DNA variants that remain polymorphic. The numbers of variants discovered using 1,000 and the whole set of 2,259 animals are marked. (**b**) Pair-wise linkage disequilibrium in four domestic breeds and wild boars from three regions. Average *r*^2^ is plotted against distance between variants. LD was calculated with common variants (MAF > 0.05) and close relatives (GRM > 0.5) removed in 435 Durocs, 522 Landraces, 493 Yorkshires, 36 Meishans, 24 European wild boars and 27 Asian wild boars. (**c**) Scatter plot of first two principal components of genotype matrix for common (MAF >0.05) and LD-pruned variants. Points are color coded according to their reported breed information. A preliminary principal component analysis was performed to visually inspect and remove clear outliers from clusters, which indicated errors in breed information. (**d**) Ancestries of pigs were estimated with variable (K = 2, 4, 6) numbers of postulated ancestral populations using the ADMIXTURE software. Estimated ancestries were plotted as stacked bar charts with breeds annotated on the top. Broad geographical locations are also annotated below the the barchart for K = 6.

Linkage disequilibrium (LD) between variants in this population was extensive but differed by breed (Figure 1b). LD in wild boars declined more rapidly as distance between variants increased than in domestic breeds, consistent with the high level of inbreeding among intensively selected domestic breeds (Figure 1b). Genetic variation present in the pig genome separated breeds into distinct clusters that represented geographic differentiation (Figure 1c,d). The first principal component of the genotypes separated Asian breeds and wild boars from their European counterparts while the second separated Durocs from other breeds (Figure 1c). Estimated ancestries of the breeds also indicated clearly separated clusters according to their geographical locations (Figure 1d). Taken together, the diverse and rich genetic variation in the 2,259 pig genomes included in this study provides a strong foundation for the development of a haplotype reference panel for imputation.

### Accuracy of genotype imputation

We focused on the approximately 34 M autosomal variants (30,489,782 SNPs and 4,125,579 indels) segregating at a minor allele frequency (MAF) > 0.005 to construct the haplotype reference panel. The distribution of MAF in the reference panel was markedly different for all variants than for those on commonly used genotyping arrays (Figure 2a). To investigate factors that influence imputation accuracy, we considered different combinations of phasing and imputation software and variable composition of the reference panel (Figure 2b). We defined imputation accuracy using three metrics, the overall concordance rate between imputed and observed genotypes, non-reference concordance rate summarizing accuracy for non-reference genotypes only, and squared correlation (*r*^2^) between imputed and observed genotypes. We evaluated three commonly used phasing/imputation software combinations and found SHAPEIT4/IMPUTE5 to outperform Beagle5.2/Beagle5.2 and Eagle2.4/Minimac4 (Figure S1) in all three metrics. SHAPEIT4/IMPUTE5 was therefore chosen for all downstream analyses.

**Figure 2.**
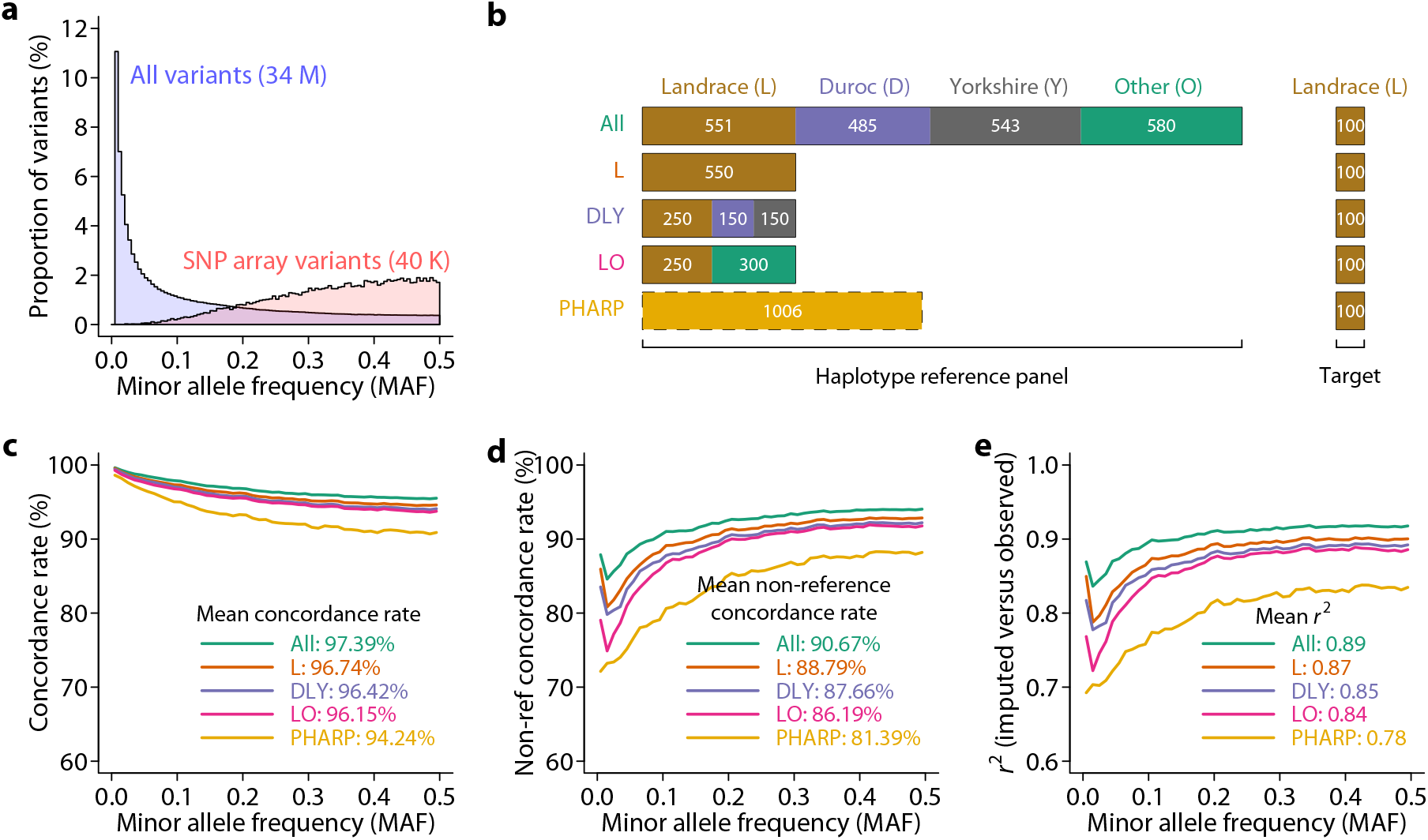
Accuracy of imputation in pigs. (**a**) Distribution of MAF for all variants in the haplotype reference panel and variants that are on the GGP Porcine 50K SNP array. Approximately 40K SNPs are present on autosomes in the reference panel. (**b**) Experimental design to investigate the effect of reference panel on imputation accuracy. Four reference panels were tested, including ‘All‘: 2,159 animals, excluding 100 Landrace pigs used as the target set; ‘L‘: 550 Landraces; ‘DLY‘: 550 pigs from Duroc, Landrace, and Yorkshire pigs; ‘LO‘: 550 pigs from Landraces and others. In addition, imputation was also performed using the PHARP server (http://alphaindex.zju.edu.cn/PHARP/index.php). (**c**) Concordance rate of imputed versus observed genotypes using different haplotype reference panels. Mean concordance rate across all variants is also indicated on the plot for each reference panel. (**d**) Non-reference concordance rate of imputed versus observed genotypes using different haplotype reference panels. Mean non-reference concordance rate across all variants is also indicated on the plot for each reference panel. (**e**) *r*^2^ of imputed versus observed genotypes using different haplotype reference panels. Mean *r*^2^ across all variants is also indicated on the plot for each reference panel.

In cattle, imputation using multi-breed reference panels appeared to be more accurate than using a single-breed panel (Daetwyler et al., 2014; Rowan et al., 2019). However, multi-breed panels are confounded by larger sample sizes. We asked whether imputation using reference panels of the same size from a single breed and from a mixture of multiple breeds made a difference (Figure 2b, compare L, DLY, and LO). This question was important as it informs whether to use a multi-breed or breed-specific reference panel to achieve optimal accuracy. We held out 100 Landrace pigs sequenced at high coverage (> 15X) and compared observed genotypes with imputed genotypes starting from genotypes at sites on a 50K SNP array. We found imputation accuracy measured by all three metrics to be remarkably similar (Figure 2c,d) when the reference panel size was equal.

Reference panel derived from the same breed as the target set had a very slight advantage (Figure 2c-e). These results suggested that the improvement in accuracy previously seen using multi-breed reference panels was likely due to the increase in sample size rather than breed composition. It is therefore determined that a multi-breed reference panel maximizing sample size is desired as opposed to breed-specific reference panel.

We compared our imputation platform using the multi-breed reference panel (*n* = 2,159, excluding the 100. Landrace test animals) with an imputation server for pigs (PHARP) (Wang et al., 2021) that utilized 1,006 animals publicly available in the SRA. We evaluated imputation accuracy among variants that were present in both reference panels. While average concordance rate was similar (97.39% versus 94.24%, Figure 2c), there was a marked improvement in non-reference concordance rate (90.67% versus 81.39%, Figure 2d) and *r*^2^ (0.89 versus 0.78, Figure 2e) using our reference panel, likely due in large part to the more than two-fold increase in sample size in this study. The final reference haplotype panel consisting of all 2,259 animals is expected to achieve concordance rate in excess of 97.39%, non-reference concordance rate of 90.67%, and *r*^2^ of 0.89.

### Genetic mapping using imputed whole sequence level genotypes

To demonstrate the usefulness of sequence level genotype imputation in genetic mapping, we performed genome-wide association studies (GWAS) for three important growth traits in pigs, using both SNP arrays and imputed genotypes.

#### Backfat thickness

Backfat thickness (BF) is one of the most important economic traits in pigs and has been intensively interrogated for its genetic basis. Genomic heritabilities estimated using either array SNPs or imputed SNPs were similar and indicated a moderately heritable trait (Figure 3a). Alleles in several genes including *IGF2* (Van Laere et al., 2003; Nezer et al., 1999), *MC4R* (Kim et al., 2000), and *LEPR* (Cristina et al., 2002) have been consistently associated with BF variation in pigs. In particular, a missense mutation in the *MC4R* gene (chr1:160773437:G>A) has been suggested as the causative mutation (Kim et al., 2000) and extensively replicated in multiple genetic backgrounds (Gozalo-Marcilla et al., 2021). Furthermore, mutations in *MC4R* are strongly associated with early onset obesity in humans (Farooqi et al., 2000) and its role in regulation of energy homeostasis is well established (Krashes et al., 2016). Importantly, the putative causal mutation in *MC4R* has been included in one of the commercially available SNP genotyping arrays, the Geneseek GGP Porcine 50K SNP Chip (Neogen, Lincoln, NE). However, the same SNP is not present in the more widely used Illumina PorcineSNP60 chip. To see if genotype imputation was able to correctly impute the genotypes of this SNP, we excluded the *MC4R* SNP and imputed whole genome genotypes from a population of 3,769 Duroc pigs genotyped using the GGP Porcine 50K SNP arrays. Remarkably, the concordance rate and *r*^2^ between the imputed and array *MC4R* SNP genotypes were 99.71% and 0.9916 respectively. Using imputed genotypes, a major peak was identified on chromosome 1, where the highest hit from imputed SNPs (chr1:161511936:T>C, *P* = 2.98 × 10^−13^) explained 2.85% of the total phenotypic variance (Figure 3a). Under this peak in a 4 Mb region (158.5 Mb – 162.5 Mb), there were 7,138 variants within 22 genes. Linkage disequilibrium in this region was extensive, with 1,050 variants in strong LD (*r*^2^ > 0.8) with the top hit, including the *MC4R* SNP (Figure 3b). The highest hit was an intronic SNP in the gene *CCBE1* (Figure 3b). However, the extensive LD in this region makes it difficult to pinpoint a causative mutation by genetic data alone. Additional functional information and genetic data that break the LD are necessary to further fine map causative genes and mutations. Nevertheless, the ability to identify the putative *MC4R* causative SNP as one of the top associated variants in a long stretch of high LD region clearly demonstrated the improvement of resolution using imputed genotypes. In our analysis, the *MC4R* SNP was initially removed and would otherwise be invisible without the imputation, as would be the case if the Illumina PorcineSNP60 chips were used.

**Figure 3.**
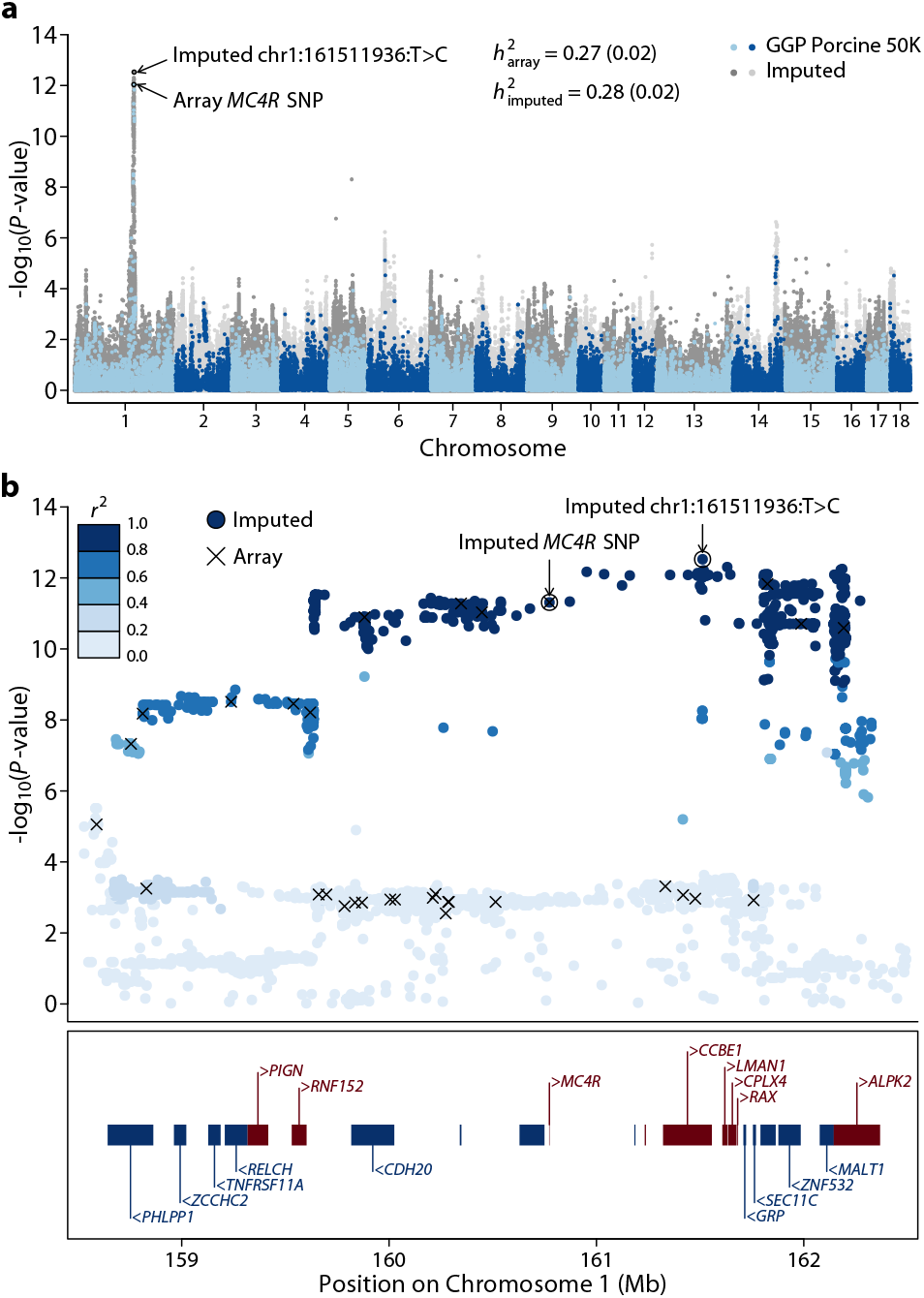
Genetic mapping of backfat thickness in pigs. (**a**) Manhattan plots of genome-wide association studies (GWAS) for backfat thickness. The grey (dark and light) points on the background are from GWAS using imputed genotypes while the blue (light and dark) points are from GWAS using SNP chips. Genomic heritabilities calculated using array and imputed genotypes are indicated. The most significant SNPs from GWAS using imputed and array genotypes are indicated by circles and arrows. (**b**) Association within the 158.5-162.5Mb region of chromosome 1, where the top hits in GWAS are located. Points indicate -log_10_(*P*-value) along the chromosome using imputed genotypes and SNPs where arrays also have genotypes are marked by crosses. The top SNPs from GWAS using imputed and array genotypes are marked by circles and arrows. *r*^2^ between the SNPs and the top SNP (chr1:161511936:T>C) are indicated by a gradient of blue color. Locations of genes are indicated in the box below the plot where blue boxes and gene names with a left arrow head (<) indicate genes transcribed on the reverse strand and red boxes and gene names with a right arrow head (>) indicate genes transcribed from the forward strand. Genes that are not marked do not have gene symbols.

#### Body length

We next considered body length. We imputed genotypes from an Affymetrix 55K SNP chip to whole genome sequence using our imputation platform and performed GWAS in a population of 1,694 Yorkshire boars (Figure 4a). The trait has a moderately high heritability, as estimated using both array (*h*^*2*^ ∼ 0.32) and imputed (*h*^*2*^ ∼ 0.34) genotypes (Figure 4a). We found a highly significant peak on chromosome 17 (Figure 4a) where the lead variant was an intergenic SNP upstream of the *BMP2* gene (chr17:15643342:C>T, *P* = 3.45 × 10^−39^). Remarkably, this variant explained 13.65% of the total phenotypic variance and the homozygous C/C animals were on average 4.01 cm longer than the T/T homozygotes (Figure 4b,c). *BMP2* has been repeatedly shown to be associated with growth traits in pigs and a recent study implicated a regulatory variant upstream of the *BMP2* gene and validated its functional impact using reporter genes (Li et al., 2021a). This regulatory variant was the third most significant SNP under this peak in our analysis.

**Figure 4.**
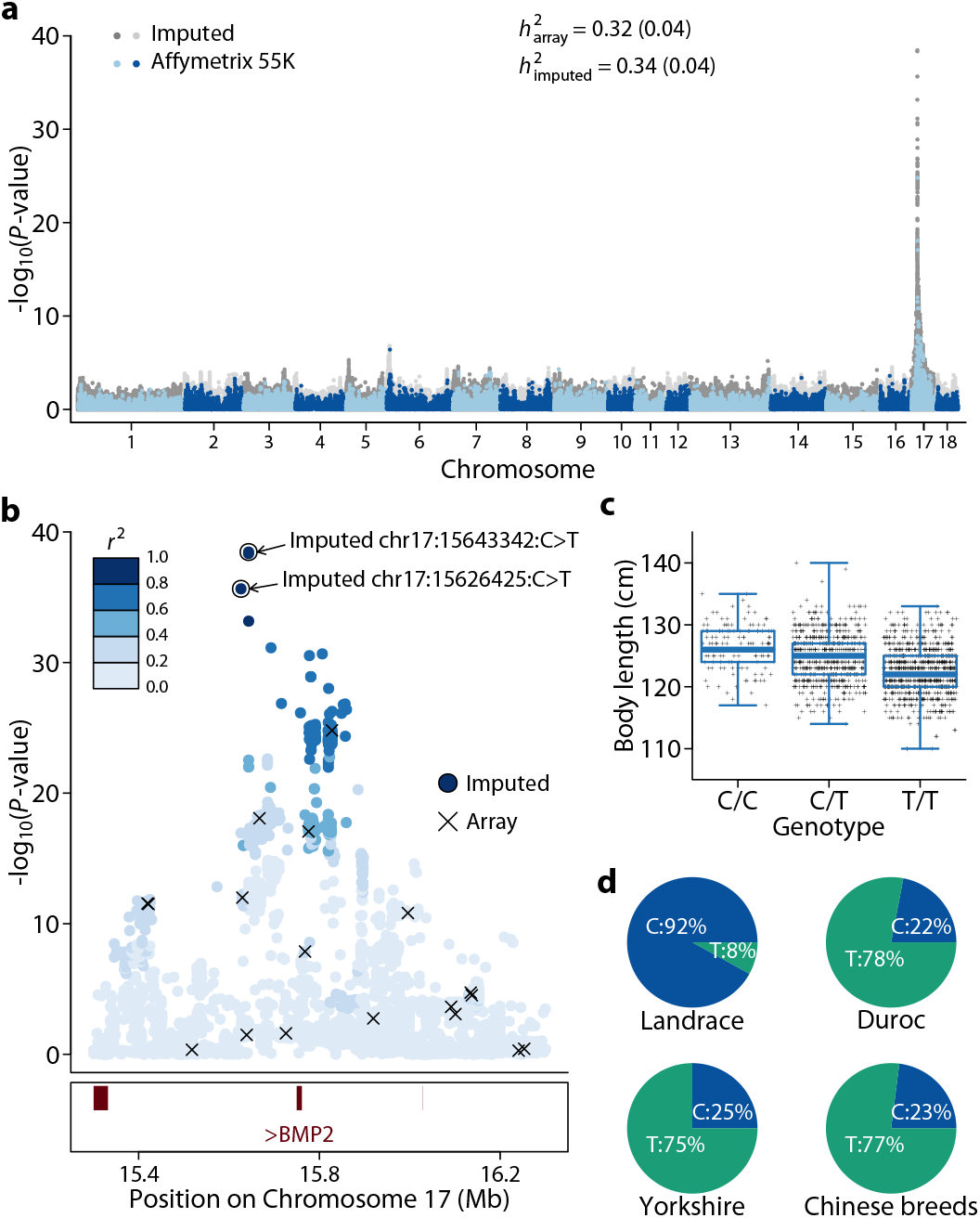
Genetic mapping of body length in pigs. (**a**) Manhattan plots of genomewide association studies (GWAS) for body length. The grey (dark and light) points on the background are from GWAS using imputed genotypes while the blue (light and dark) points are from GWAS using SNP chips. Genomic heritabilities calculated using array and imputed genotypes are indicated. (**b**) Association within the 15.3-16.3Mb region of chromosome 17, where the top hits in GWAS are located. Points indicate -log_10_(*P*-value) along the chromosome using imputed genotypes and SNPs where arrays also have genotypes are marked by crosses. The top SNPs from GWAS using imputed and array genotypes are marked by circles and arrows. *r*^2^ between the SNPs and the top SNP (chr17:15643342:C>T) are indicated by a gradient of blue color. Locations of genes are indicated in the box below the plot. All three genes are colored in red and transcribed from the forward strand. The only gene with a symbol in this region is BMP2. (**c**) Scatter and box plots of body length (in cm) for the three genotypes of the chr17:15643342:C>T SNP. (**d**) Allele frequencies of the chr17:15643342:C>T SNP in different breeds.

Whether one or both of these potentially regulatory variants are the causative mutations remain to be determined. Given the strong association, high MAF of these SNPs, and less extensive LD in this region, it is unlikely that these regulatory variants were tagging protein-coding and less common variants in the BMP2 gene. In addition to the genetic support from this Yorkshire population, the body length increasing C allele was much more prevalent in Landrace than in other breeds. A hallmark of the Landrace breed is its long body size thus regulatory variation of the *BMP2* gene may be a major contributor to the phenotypic differentiation between pig breeds. In contrast, although the SNP chip was able to broadly identify this region, the most significant SNP (chr17:15827832:T>G, *P* = 1.58 × 10^−25^) in a SNP chip based GWAS was about 184 kb away from the lead SNP and explained a substantially smaller variance (8.22% versus 13.65%).

#### Birth weight

Lastly, we mapped genes for birth weight, a trait with low heritability (*h*^2^ ∼ 0.14, Figure 5a) and few known mapped genes. No SNP was above a genome-wide significance threshold (*P* = 5 × 10^−8^) using SNP chips in a population of 2,855 Duroc pigs. In contrast, a significant peak was identified under a short insertion (chr2:7619832:C>CCT, *P* = 3.86 × 10^−8^), which explained 1.61% of phenotypic variation in birth weight (Figure 5a). The indel is located upstream of the *SLC22A11* gene (Figure 5b), which is one of the members in the SLC22 transporter family. *SLC22A11* in humans is expressed in placenta, kidney, and brain and transports metabolites such as urate and estrone sulfate (Nigam, 2018). *SLC22A11* plays an important role in the uptake of the fetal liver derived 6*α*-hydroxydehydroepiandrosterone sulfate from to placental syncytiotrophoblasts during human pregnancy (Tomi et al., 2015). While its role isn’t definitive, this example represents a scenario where imputation identified candidate genes that were completely missed by GWAS using SNP chips.

**Figure 5.**
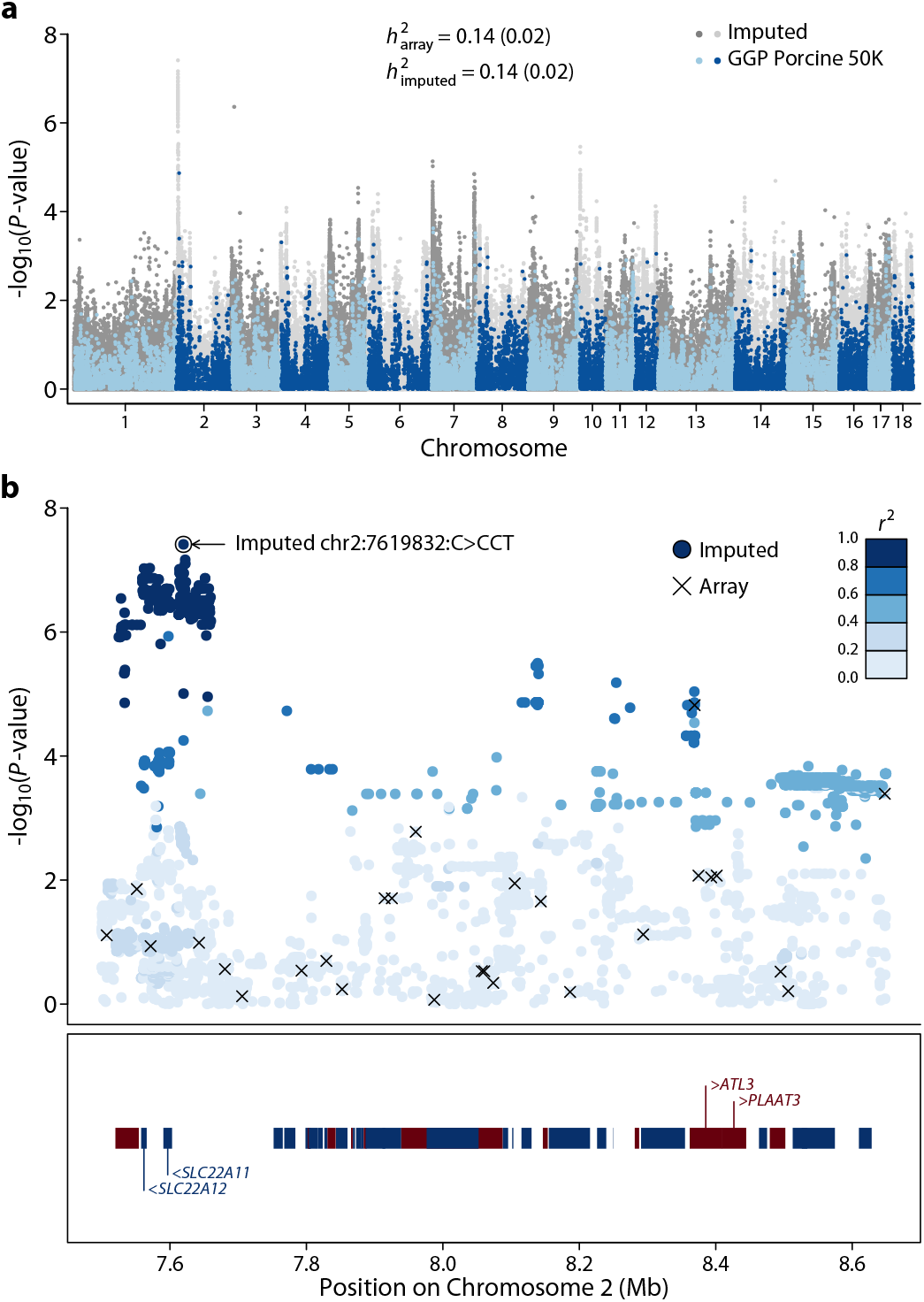
Genetic mapping of birth weight in pigs. (**a**) Manhattan plots of genomewide association studies (GWAS) for birth weight. The grey (dark and light) points on the background are from GWAS using imputed genotypes while the blue (light and dark) points are from GWAS using SNP chips. Genomic heritabilities calculated using array and imputed genotypes are indicated. (**b**) Association within the 7.50-8.65Mb region of chromosome 2, where the top hits in GWAS are located. Points indicate -log_10_(*P*-value) along the chromosome using imputed genotypes and SNPs where arrays also have genotypes are marked by crosses. The lead SNP from GWAS using imputed genotypes is marked by a circle and an arrow. *r*^2^ between the SNPs and the top SNP (chr2:7619832:C>CCT) are indicated by a gradient of blue color. Locations of genes are indicated in the box below the plot. The region contains many genes, and only genes near the top GWAS hits are indicated.

### Swine Imputation (SWIM) server

To enable the broad research community to efficiently utilize the resource developed in this study, we developed a public SWine IMputation (SWIM) web server (https://www.swimgeno.org), on which users can upload SNP chip genotypes and retrieve imputed genotypes. The user interface is extremely simple, which only requires users to upload the genotypes in gzipped ped/map format and leave their emails. Allele matching and flipping are performed on the server end, further simplifying the process on the user end. Imputation status can be monitored and results can be downloaded from a dynamic link without having to register an account. The server is set up to accommodate multiple users at the same time while limiting multiple jobs of the same user (can still be queued). Our tests indicated that a typical job with 2,000 individuals and 50K SNP chip genotypes can be completed in approximately 12 hours.

## Discussion

We present here the development of the largest reference haplotype panel in pigs and an accompanying web server for the public to utilize this resource for genotype imputation. The high level of diversity and the large number of animals in the panel enabled us to achieve very high imputation accuracy with concordance rate, non-reference concordance rate, and *r*^2^ in excess of 97.39%, 90.67%, and 0.89 respectively (Figure 2). Given the high accuracy and easy access, we expect this public resource to vastly democratize sequence level imputation in pigs and accelerate genetic discoveries.

High throughput genotyping arrays greatly simplified genotyping and numerous new QTLs have been mapped by association mapping, typically within a breed and with hundreds to thousands of individuals (Hu et al., 2019). However, while resolution has improved with SNP arrays, causative genes and mutations remain extremely elusive, partly because SNP arrays prioritize assay feasibility, homogeneous spacing, and common SNPs (Ramos et al., 2009).

As we have shown with the examples above, imputation is expected to greatly improve resolution of gene mapping. Given the large number of existing genome-wide association studies in pigs (Hu et al., 2019), we expect this resource to be highly utilized and impactful. All existing studies using SNP arrays can be improved by a simple imputation followed by GWAS without additional data. Metaanalysis also becomes possible because a common SNP set can be obtained. Nonetheless, the resolution of genetic mapping depends not only on SNP density but also on experimental design and genetic structure in the mapping population. Sequence level imputation does not necessarily identify causative mutations in one single step (Yan et al., 2022). The availability of this resource will allow for suitable designs of mapping studies to achieve the highest possible resolution in specific circumstances and potentially nucleotide resolution.

## Materials and Methods

### WGS data collection

We consolidated WGS data from multiple sources. A total of 1,530 animals are first reported in this study using Illumina (*n* = 863) and BGI (*n* = 667) platforms with 150 bp paired-end reads. Among them, 610 Landrace, 413 Duroc, 391 Yorkshire, 18 Taiwanhei and 17 Lichahei were from Wen’s Food Group Co., Ltd. (Yunfu, Guangdong, China), 21 Dahuabai, 21 Lantanghei, 20 Guangdong Xiaoerhua and 19 Yuedonghei from Guangdong Gene Bank of Livestock and Poultry (Guangzhou, Guangdong, China). Additionally, sequences for 729 animals were downloaded from the sequence read archive (SRA). A complete breakdown including accession numbers, sample sizes, and average sequencing coverage can be found in Table S1-2.

### Variant calling, recalibration and filtering

We aligned sequence reads to the pig reference genome (Sscrofa11.1, a Duroc pig) (Warr et al., 2020) using BWA-MEM-0.7.17 (Li and Durbin, 2009) and called variants (in GVCF format) using GATK-4.1.8.1 HaplotypeCaller (DePristo et al., 2011) after several post-alignment processing steps including duplicate removal using PicardTools-2.23.3 (DePristo et al., 2011), and base quality recalibration using GATK. A population VCF was generated by combining GVCFs across all samples. Variants with excessive heterozygosity (“ExcessHet > 54.69”) were removed. Variant quality score recalibration (VQSR) on SNPs was performed with truth SNP sets compiled from commercial SNP arrays including 50K, 60K, 80K SNP chips (prior = 15.0) on the Illumina platform and the 660K (prior = 12.0), SowPro90 (prior = 15.0) SNP chips from the Affymetrix platform. SNPs were filtered with a truth sensitivity filter level at 99.0. Without a truth set of indels, we applied hard filtering on by excluding indels with QD < 2.0 QUAL < 50.0 FS > 100.0 ReadPosRankSum < -20.0, as recommended by GATK’s best practices. Additionally, we filtered out animals with missing rate > 0.20, heterozygosity > 0.20, and retained bi-allelic sites with missing rate < 0.2, and mean sequencing depth between 5 and 500. Filtering was performed using a combination of VCFtools 0.1.13 (Danecek et al., 2011) and BCFtools 1.13 (Danecek et al., 2021) commands.

### Population genetics analysis

Linkage disequilibrium (Figure 1b) was computed using PopLDdecay (Zhang et al., 2019) on individuals within the same breed with close relatives (GRM > 0.5) and low frequency variants (MAF < 0.05) removed. To understand the genetic structure in the population, we retained variants with MAF > 0.05 and missing rate < 0.1 and pruned SNPs with LD (*r*^2^ < 0.3, -indep-pairwise 50 10 0.3) using PLINK 1.9 (Chang et al., 2015). Principal component analysis (PCA) was performed on the filtered list of 1,223,882 variants using GCTA 1.93.2 (Yang et al., 2011) for all individuals. Ancestries were estimated using ADMIXTURE 1.3 (Alexander et al., 2009) on 185 individuals randomly selected according to breed representation in the dataset or at least 4 individuals per breed. The downsampling was necessary to properly visualize population structure.

### Genotype imputation

We further filtered variants prior to phasing haplotypes in the reference population. Variants with missing rate > 0.1 and MAF < 0.005 were removed. Additionally, variants with a Hardy-Weinberg equilibrium test *P* value < 10^−10^ implemented separately in PLINK in all three of the Duroc, Landrace, and Yorkshire pigs were removed. Only autosomal variants were retained for imputation.

We extracted 100 Landrace pigs with the highest sequencing depth (17.42 X average sequencing depth, ranging from 14.98 to 63.11 X) and designated these individuals as the target population to evaluate imputation accuracy. To test the effect of breed composition of the reference population, we constructed four reference haplotype panels using different sets of individuals, including All (*n* = 2,159): all individuals except the 100 Landraces; L (*n* = 550): Landrace pigs only; DLY (*n* = 550): 250 Landraces + 150 Durocs + 150 Yorkshires; and LO (*n* = 550): 250 Landraces + 300 randomly selected pigs other than Durocs and Yorkshires. Phasing was independently performed in these reference sets. In addition, we also tested imputation using the PHARP web server (http://alphaindex.zju.edu.cn/PHARP/index.php), which contains reference haplotypes constructed from 1,006 individuals in the SRA.

We tested three combinations of software for phasing and imputation, including SHAPEIT 4.2 (Delaneau et al., 2019) + IMPUTE5 1.1.5 (Rubinacci et al., 2020), Beagle 5.2 (Browning et al., 2021) + Beagle 5.2, and Eagle 2.4 (Loh et al., 2016) + Minimac 4 (Howie et al., 2012). All software was run with default options and uninformative linkage map (1cM per 1Mb) but effective population size set to 100. Imputed genotypes were called by the ones with the highest posterior genotype probability. However, users of the imputation web server also receive genotype probabilities.

We considered three commonly used metrics of imputation accuracy, concordance rate, nonreference concordance rate (Li et al., 2021b) and *r*^2^. Concordance rate is defined as the proportion of individuals with imputed genotypes in concordance with observed genotypes. Non-reference concordance rate is similar to concordance rate but is restricted to only individuals that are not homozygous for the reference allele. *r*^2^ is the squared Pearson correlation coefficient between observed and imputed genotypes. We measured concordance rates and *r*^2^ on a per SNP basis and averaged them over SNPs in MAF bins (0.01 in size) or across the whole genome.

### Genotypic and phenotypic data collection

To demonstrate the utility of imputation in genetic mapping, we collected phenotypes and genotypes for three populations of pigs, which were managed by three core breeding farms of Wen’s Food Group Co., Ltd. (Yunfu, Guangdong, China), all under standard management practices. For backfat thickness, the phenotypes were collected on 3,769 Duroc pigs from 2013 to 2018, and SNP genotyping was performed using the Geneseek GGP Porcine 50 K SNP chip (Neogen, Lincoln, NE, USA). Backfat thickness was measured between the 10th and 11th ribs using an Aloka 500V SSD B ultrasound (Corometrics Medical Systems, USA) when live weights of pigs reached about 100 kg (100 ± 5 kg). For body length, phenotypes from a total of 1,694 Yorkshire boars were collected from 2012 to 2018, and SNP genotyping was performed using the Affymetrix PorcineWens55K SNP chip (Affymetrix, Santa Clara, CA, United States). Body length was measured from the base of the ear to the base of the tail in pigs at approximately 100 kg (100 ± 5 kg) body weight. For birth weight, a total of 2,857 Duroc pigs were phenotyped from 2015 to 2018, and SNP genotyping was performed using the Geneseek GGP Porcine 50 K SNP chips. All samples were collected according to the guidelines for the care and use of experimental animals approved by the Ministry of Agriculture and Rural Affairs of the People’s Republic of China. The ethics committee of South China Agricultural University specifically approved the animal use in this study.

### Genome-wide association studies

We used GCTA 1.92.1 to perform a mixed linear model (MLM) based association analysis. The following statistical model was used: *y* = *µ* + *xb* + *g* + *e*, where y is the vector of the phenotypic values for all animals, *µ* is the con, *x* is the design matrix coding genotypes and other incidences of fixed effects, *b* is the vector of fixed effects including SNP effect and additional covariates such as sex, pen, year-season effects depending on the traits and, *g* is the vector of polygenic random effects with covariance dictated by the genomic relationship matrix, and *e* is the vector of random residuals. We used a genome-wide significance threshold of *P* = 5 × 10^−8^ to declare significance.

## Data availability

Raw sequence data for a subset of the animals utilized in this study were downloaded from SRA (Table S1-2). Additional sequenced animals were proprietary properties of Wen’s Food Group Co., Ltd. and Guangdong Gene Bank of Livestock and Poultry. They may be requested by contacting respectively. Imputation utilizing the full data set is delivered as a web service (www.swimgeno.org) and is publicly available.

## Code availability

All computer codes including all analyses performed in this study and codes for the SWIM web server are available at https://github.com/qgg-lab/swim-public.

## Acknowledgement

This work is supported by a USDA-NIFA project (2021-67021-34149, to W.H., C.G., J.S., R.S), a USDA-NIFA Hatch project (MICL 02560, to W.H.), a Natural Science Foundation of China project (31972540, to J.Y.), and a Natural Science Foundation of Guangdong Province project (2018B030313011, to Z.W.). The web server is supported by the USDA Swine Genome Coordinator Fund (NRSP8).

## Conflict of Interest

C.T. and G.C. are employees of the Guangdong Zhongxin Breeding Technology Co., Ltd.

