## Supplemental figures for "Nucleotide resolution genetic mapping in pigs by publicly accessible whole genome imputation"

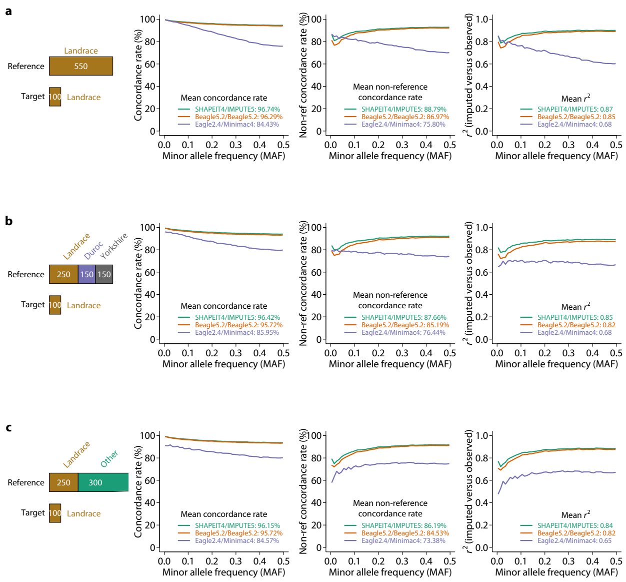


**Figure S1 | Comparison of software combinations for imputation.** (**a**) Concordance rate, non-reference concordance rate, and $r^2$ of imputed versus observed genotypes using different software combinations with 550 Landraces as the reference panel. (**b**) Same analysis but in a reference panel consisting of 250 Landraces, 150 Durocs, and 150 Yorkshires. (**c**) Same analysis but in a reference panel consisting of 250 Landraces and 300 other breeds (not Duroc or Yorkshire).
